# Identification of 12 genetic loci associated with human healthspan

**DOI:** 10.1101/300889

**Authors:** Aleksandr Zenin, Yakov Tsepilov, Sodbo Sharapov, Evgeny Getmantsev, L. I. Menshikov, Peter O. Fedichev, Yurii Aulchenko

**Affiliations:** Gero LLC, Novokuznetskaya street 24/2, Moscow 119017, Russia; Novosibirsk State University, Pirogova 2, 630090, Novosibirsk, Russia; Institute of Cytology and Genetics SB RAS, Lavrentyeva ave. 10, 630090, Novosibirsk, Russia; National Research Center Kurchatov Institute, 1, Akademika Kurchatova pl., Moscow 123182, Russia; Moscow Institute of Physics and Technology, 141700, Institutskii per. 9, Dolgoprudny, Moscow Region, Russia; PolyOmica, Het Vlaggeschip 61, 5237PA, ’s-Hertogenbosch, The Netherlands

## Abstract

The mounting challenge of preserving the quality of life in an aging population directs the focus of longevity science to the regulatory pathways controlling healthspan. To understand the nature of the relationship between the healthspan and lifespan and uncover the genetic architecture of the two phenotypes, we studied the incidence of major age-related diseases in the UK Biobank (UKB) cohort. We observed that the incidence rates of major chronic diseases increase exponentially. The risk of disease acquisition doubled approximately every eight years, i.e., at a rate compatible with the doubling time of the Gompertz mortality law. Assuming that aging is the single underlying factor behind the morbidity rates dynamics, we built a proportional hazards model to predict the risks of the diseases and therefore the age corresponding to the end of healthspan of an individual depending on their age, gender, and the genetic background. We suggested a computationally efficient procedure for the determination of the effect size and statistical significance of individual gene variants associations with healthspan in a form suitable for a Genome-Wide Association Studies (GWAS). Using the UKB sub-population of 300,447 genetically Caucasian, British individuals as a discovery cohort, we identified 12 loci associated with healthspan and reaching the whole-genome level of significance. We observed strong (|*ρ_g_*| > 0.3) genetic correlations between healthspan and the incidence of specific age-related disease present in our healthspan definition (with the notable exception of dementia). Other examples included all-cause mortality (as derived from parental survival, with *ρ_g_* = −0.76), life-history traits (metrics of obesity, age at first birth), levels of different metabolites (lipids, amino acids, glycemic traits), and psychological traits (smoking behaviour, cognitive performance, depressive symptoms, insomnia). We conclude by noting that the healthspan phenotype, suggested and characterized here, offers a promising new way to investigate human longevity by exploiting the data from genetic and clinical data on living individuals.

## I. INTRODUCTION

Age is the most important single risk factor for multiple diseases, see, e.g., [1]. Likewise, extreme longevity in human cohorts is associated with a delayed incidence of diseases: KaplanMeyer curves of disease-free survival, stratified by age, demonstrate a consistent delay in the onset of age-related diseases with increasing age of survival [2]. The emerging premise is, therefore, that aging itself is the common driver of chronic diseases and conditions that limit the functional and disease-free period, the healthspan, and hence is a target for novel interventions [3]. The increasing number of available genomes of very old people [4–6], though representing a rather specific and a relatively small sub-group of exceptionally successfully aging individuals, can provide an insight on the genetic architecture of exceptional life- and health-spans by use Genome-Wide Association Studies (GWAS). While such studies suggested a fair number of loci, the *APOE/TOMM40* locus is probably among the few consistently implicated in multiple studies. Further gains can be naturally achieved by increasing the population size with the help of proxy phenotypes, such as a search for genetic variants that predispose one to age-related disease and hence are depleted in long-lived persons compared to controls [5]. Another promising alternative involves GWAS of parental lifespans [7–9].

In this paper, we focused on aging and morbidity in mid-life and introduce human healthspan as an alternative phenotype for genetics of longevity studies. More specifically, we used clinical histories for over 300,000 people, the participants of the UK Biobank (UKB) cohort in the age range from 37 to 72 years old. We checked the incidence of chronic diseases and identified the top eight morbidities strongly associated with age after the age of 40 and ranked by the number of occurrences in the UKB cohort. We observed that the risk of the selected diseases increases exponentially at approximately the same rate. The corresponding doubling time is approximately eight years, close to the mortality risk doubling time from Gompertz law of mortality [10]. The close association between disease and mortality risk dynamics suggested the possibility of a single underlying mechanism, that is aging.

To reveal the genetic determinants of healthspan, we built a proportional hazards model to predict the end of the disease-free period of an individual depending on their age, gender, and the genetic background. We used the sub-population of 300, 447 genetically Caucasian individuals as a discovery cohort for a GWAS and identified 12 loci associated with healthspan at the whole-genome level of significance. The genetic signature of healthspan has high genetic correlations with GWAS of obesity, type 2 diabetes, coronary heart disease, traits related to metabolic syndrome, and all-cause mortality (as derived from parental survival). We conclude by noting that the healthspan phenotype, proposed and characterized here, offers a promising new way to investigate human longevity by exploiting the data from large cohorts of living individuals with rich clinical information.

## II. RESULTS

### A. Healthspan and longevity in UKB cohort

We studied the dynamics of disease incidence using the clinical data available from UKB and selected the top eight morbidities strongly associated with age after the age of 40 and ranked by the number of occurrences. The shortlist included Congestive Heart Failure (CHF), Myocardial Infarction (MI), Chronic Obstructive Pulmonary Disease (COPD), stroke, dementia, diabetes, cancer, and death (Table S11). The risks of the selected conditions were found to increase exponentially with age at approximately the same rates (Fig. 1; see Materials and Methods section A4 for details). The characteristic doubling time is approximately seven to eight years. The risk of death in the dataset also grows exponentially with age following empirical Gompertz mortality law [10, 11]. The manifested similarity between the diseases and the mortality risk doubling time suggest that the most plausable single unifying mechanism behind the risk acceleration with age is aging itself.

**FIG. 1:**
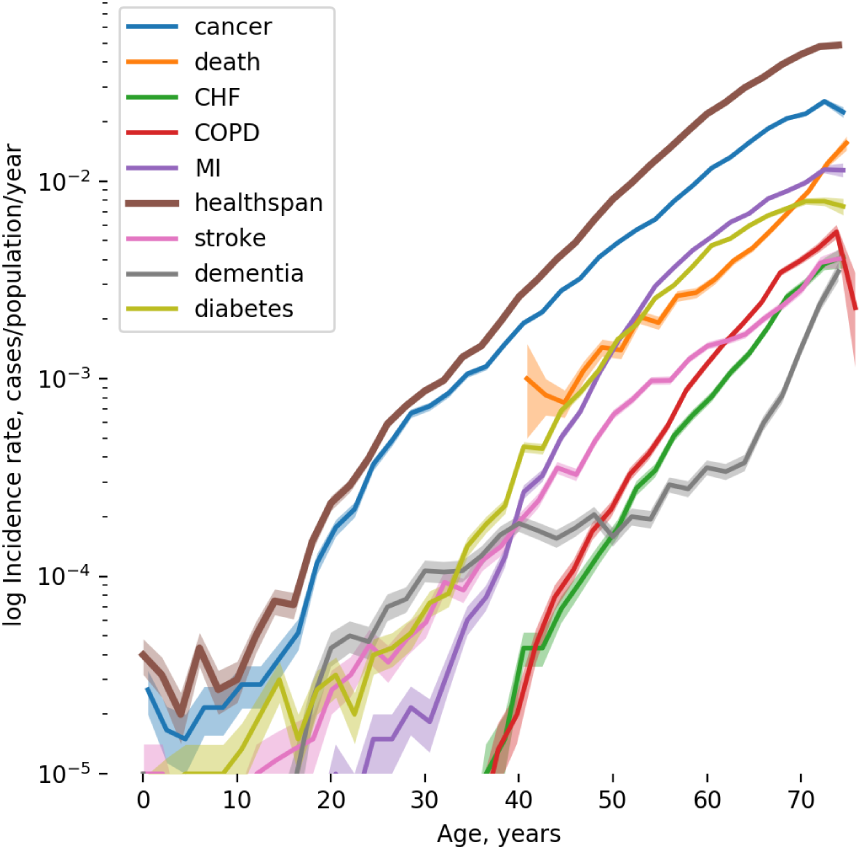
The incidence of the most prevalent chronic diseases (the solid lines) and the risk of death (the mortality rate, the orange line) for UKB participants. The disease incidence increases approximately exponentially with age at approximately the same rates.

We chose to define healthspan as the age of the onset of the first disease from our predefined list of age-dependent diseases. As expected, the first morbidity incidence rate also increases exponentially with age (see the brown “healthspan” line in Fig. 1), the corresponding doubling time matches the mortality, and the specific disease risk doubling times. In the UKB cohort, healthspan is ended by cancer in more than half of the cases, followed by diabetes and MI, see Figure 2. These three diseases alone account for over 86% of the end of healthspan period (although the cancer category itself includes a huge variety of diseases). Death occurs later in life and follows the end of the disease-free survival by approximately a decade. The total number of the participants with one or more chronic diseases, 84, 949, is dramatically larger than that of death events, 8, 365, out of 300, 447 study population (see below for the GWAS inclusion criteria).

**FIG. 2:**
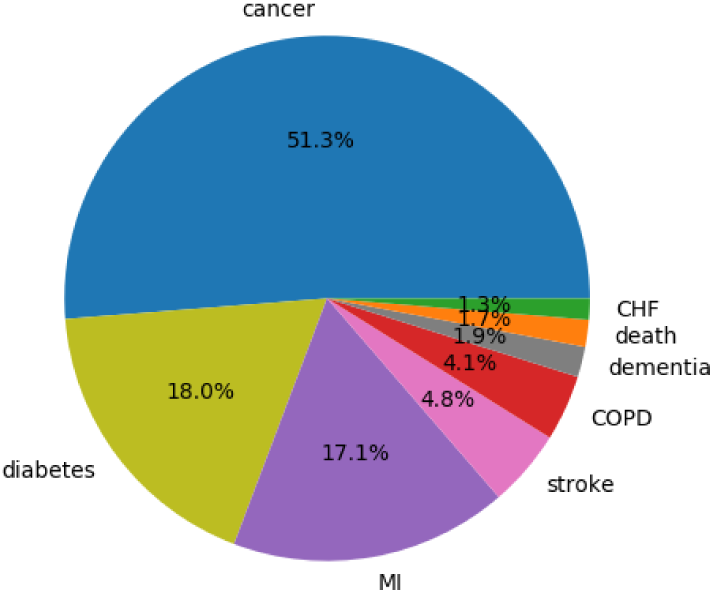
Pie chart representing the first chronic disease prevalence from UKB clinical information (the diseases color codes are the same as in Figure 1).

### B. Genome-wide association study design

Next, we identified gene-variants predisposing individuals to a shorter healthspan. Since the incidence of the first morbidity risk grows exponentially with age, we propose to employ the Cox-Gompertz proportional model (see, e.g., [12]) to test statistical associations between specific genes and disease risks. In Appendix A 5 we explain how to use a maximum likelihood version of Cox-Gompertz model to predict the age corresponding to the end of healthspan for each study participant.

We started by characterizing each of the 300,447 individuals in the study cohort by sex and age, followed by the technical (genotyping batch, assessment center), and the ethnicity-related genetic variables (40 first genetic principal components). Maximum likelihood optimization produced the best fit proportional hazards model parameters. The morbidity incidence growth rate was found to be 0.098 per year, which corresponds to a doubling time of seven years and is compatible with the mortality rate doubling time of approximately eight years from Gompertz mortality law. As expected, being male is a significant risk factor (log-hazard ratio, log(*HR*) = 0.26 at the significance level of *p* < 10^−10^), with a corresponding healthspan difference of approximately three years. The genetic principal components PC4 and PC5, and some of the assessment center labels are also highly significantly associated with the healthspan. From these numbers, we observed that human mortality and the first morbidity incidence follow a version of Gompertz law. The average healthspan can be readily estimated from the Gompertz model parameters as 72 years, which is 14 years less than the Cox-Gompertz lifespan estimate for the same cohort.

Since we did not expect a substantial effect on healthspan by any of the individual gene-variants, the effect sizes and the significance testing could be performed using a form of linear regression to the Martingale residual of the Cox-Gompertz model above, see Appendix A 6. In this study, we limited the discovery association screen to the study cohort (300, 447 individuals) with available genetic information with 11, 309, 218 imputed autosomal variants.

### C. GWAS results

A total of 394 SNPs at 14 loci achieved a genome-wide significance threshold of *p* < 5 × 10^−8^ (Table S9). Using median estimator, the genomic control inflation parameter λ [13] was 1.18. The LD score regression [14] yielded the healthspan heritability of 0.102 (*se* = 0.009), and the LD score regression intercept was 1.053 (*se* = 0.008, *ratio* = 0.24). After adjusting the results of the discovery GWAS for genomic control of 1.053, a total of 328 SNPs positioned in twelve loci remained statistically significant at the genome-wide level (Fig. 3). The conditional and joint analysis confirmed that all the regions were independent except a locus on chromosome 6, at 161Mb (Table SI). We detected two signals in this locus (rs14057088 and rs10455872) that were in very weak linkage disequilibrium (*R* = −0.04). Interestingly, the distance between these SNPs was only 3kbp, and they had relatively small frequencies (0. 08 and 0.016, respectively).

**FIG. 3:**
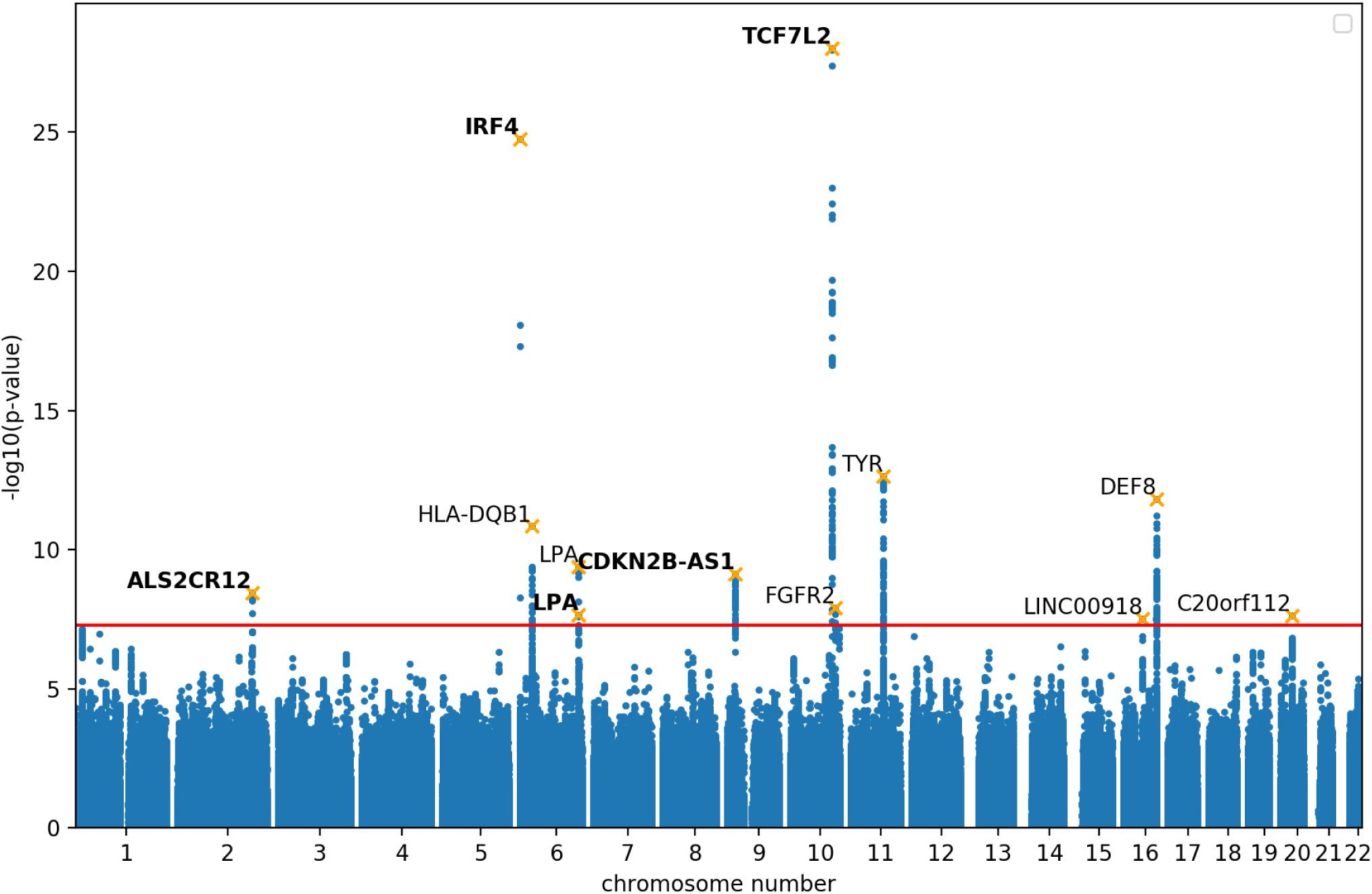
Manhattan plot representing the discovery analysis of healthspan in genetically Caucasian individuals. The trait is a form of Martingale residual of the Cox-Gompertz proportional hazards model of healthspan as described in Section A 5. The loci are tagged by SNPs from Table I, labeled by the nearest gene symbol, replicated SNPs marked in bold.

**TABLE I:**
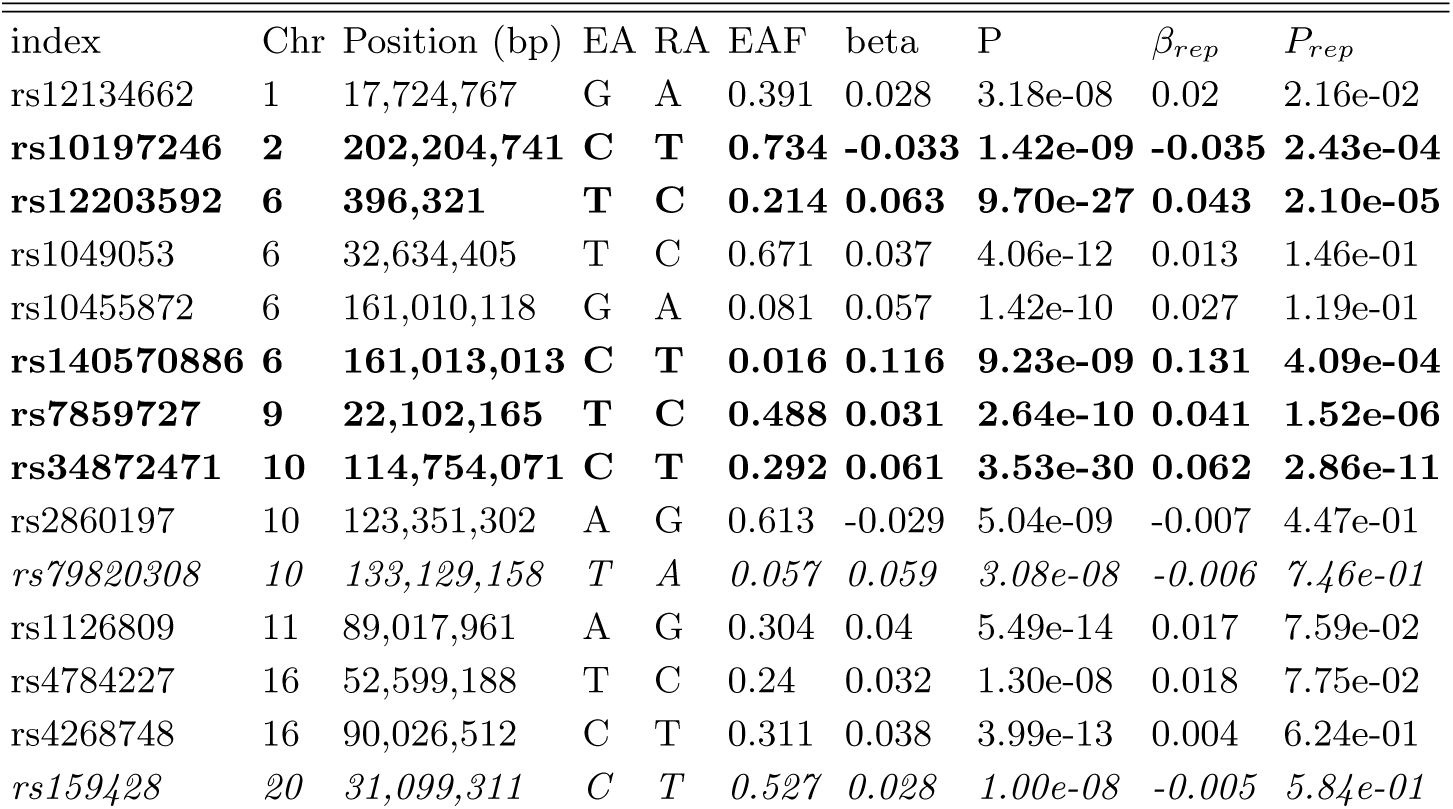
Variants, tagging regions, significantly associated with the first morbidity hazard (end of healthspan) in 300,447 genetically British individuals, and results of replication in 96,313 individuals. EA: effective (coded, tested) allele, RA: reference (non-coded) allele, EAF: effect allele frequency, *β*: regression coefficient estimate, p: p-value, *β_rep_*: regression coefficient estimate in replication sample, *p_rep_*: p-value in replication sample. In bold: replicated loci. In italics: loci demonstrating opposite effect in replication.

| index | Chr | Position (bp) | EA | RA | EAF | beta | P | $\beta_{rep}$ | $P_{rep}$ |
| --- | --- | --- | --- | --- | --- | --- | --- | --- | --- |
| rs12134662 | 1 | 17,724,767 | G | A | 0.391 | 0.028 | 3.18e-08 | 0.02 | 2.16e-02 |
| **rs10197246** | **2** | **202,204,741** | **C** | **T** | **0.734** | **-0.033** | **1.42e-09** | **-0.035** | **2.43e-04** |
| **rs12203592** | **6** | **396,321** | **T** | **C** | **0.214** | **0.063** | **9.70e-27** | **0.043** | **2.10e-05** |
| rs1049053 | 6 | 32,634,405 | T | C | 0.671 | 0.037 | 4.06e-12 | 0.013 | 1.46e-01 |
| rs10455872 | 6 | 161,010,118 | G | A | 0.081 | 0.057 | 1.42e-10 | 0.027 | 1.19e-01 |
| **rs140570886** | **6** | **161,013,013** | **C** | **T** | **0.016** | **0.116** | **9.23e-09** | **0.131** | **4.09e-04** |
| **rs7859727** | **9** | **22,102,165** | **T** | **C** | **0.488** | **0.031** | **2.64e-10** | **0.041** | **1.52e-06** |
| **rs34872471** | **10** | **114,754,071** | **C** | **T** | **0.292** | **0.061** | **3.53e-30** | **0.062** | **2.86e-11** |
| rs2860197 | 10 | 123,351,302 | A | G | 0.613 | -0.029 | 5.04e-09 | -0.007 | 4.47e-01 |
| *rs79820308* | *10* | *133,129,158* | *T* | *A* | *0.057* | *0.059* | *3.08e-08* | *-0.006* | *7.46e-01* |
| rs1126809 | 11 | 89,017,961 | A | G | 0.304 | 0.04 | 5.49e-14 | 0.017 | 7.59e-02 |
| rs4784227 | 16 | 52,599,188 | T | C | 0.24 | 0.032 | 1.30e-08 | 0.018 | 7.75e-02 |
| rs4268748 | 16 | 90,026,512 | C | T | 0.311 | 0.038 | 3.99e-13 | 0.004 | 6.24e-01 |
| *rs159428* | *20* | *31,099,311* | *C* | *T* | *0.527* | *0.028* | *1.00e-08* | *-0.005* | *5.84e-01* |

Using meta-analysis for the subsets of the UK Biobank that we selected for replication (total *N* = 96,313), we performed replication of 12 genome-wide significant SNPs (Table SI). Of the twelve SNPs, all but one demonstrated a consistent sign of association in replication. Of the eleven SNPs with consistent signs, five associations were significant after correction for multiple testing with *p* < (0.05/12). We subsequently refer to these SNPs as ‘replicated’.

### D. Genetic correlation analysis

First we checked the genetic correlations between the healthspan GWAS results and the genetic signatures of the individual diseases used to build the healthspan phenotype. To do this, we produced a series of independent GWAS of the age at onset of the individual conditions, using the same Cox-Gompertz methodology (Fig. 4). The healthspan GWAS exhibits strong correlations with most of the disease traits, with the notable exception of dementia (see the discussion below). What is more assuring, the mortality, stroke, CHF and MI traits yield better correlations with healthspan than cancer, even though cancer is the most frequently occurring first morbidity leading to termination of the disease free period in our study. We therefore conclude that most of the selected diseases (perhaps, with the exception of the dementia) and all-cause mortality share a significant number of common genetic determinants.

To obtain a broader insight into biological significance of our findings we analyzed genetic correlations between healthspan and 235 complex traits available from the LD-hub [15]. Overall, we observed significant genetic correlations (*p* < 4.3 × 10^−5^) between the frailty risk and 46 traits (Table S2). The strongest positive correlations (*r_g_* > 0.4) were found in association with coronary artery disease (CAD) [16] (*r_g_* = 0.62), Type 2 Diabetes [17] (*r_g_* = 0.58), glycated hemoglobin level (HbA1C) [18] (*r_g_* = 0.42), cigarettes smoked per day [19] (*r_g_* = 0.44), and insulin resistance index (HOMA-IR) [20] (*r_g_* = 0.41). The strongest negative correlations (*r_g_* < −0.4) were for the age of first birth [21] (*r_g_* = −0.43), father’s, mother’s, parental age at death [22] (*r_g_* = −0.74, −0.66, −0.76 respectively), former vs. current smoker [19] (*r_g_* = −0.48) and HDL related traits [23] (cholesterol esters in large HDL, total lipids in large HDL, total cholesterol in large HDL, mean diameter for HDL particles, free cholesterol in large HDL, with *r_g_* = −0.44, −0.41, −0.44, −0.42 and −0.43, respectively).

**FIG. 4:**
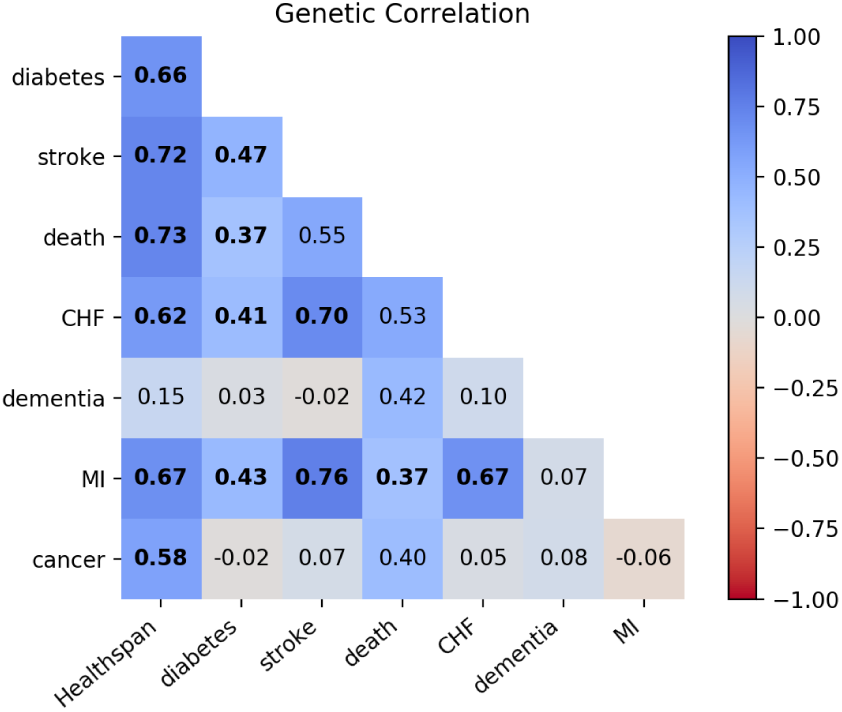
Genetic correlation between GWAS of the healthspan and the diseases used to produce the healthspan phenotype in the UKB discovery cohort. The significant correlations marked in bold (*p* < 0.05 after Bonferroni correction). The numbers corresponding to the COPD trait are not shown, since the GWAS fails to produce summary statistics due to mean Z score above 0.15.

Figure 5 summarizes the results of clustering analysis of the top genetic correlations selected by significance. We found, that 35 traits with strong significant genetic correlation with healthspan (|*r_g_*| > 0.3 and *p* < 4.3 × 10^−5^) fall into four distinct clusters: 1) the group of sociodemographic factors (including education), lifespan traits, smoking, CAD and lung cancer; 2) HDL-related traits; 3) the cluster of obesity-related traits including BMI and 4) Type 2 diabetes-related traits. The healthspan itself clusters together with CAD and parental lifespan (a sub-cluster of cluster 1). We note, however, the absence of any substantial genetic correlation between the healthspan and Alzheimer disease (*r_g_* = −0.03, Table S2).

**FIG. 5:**
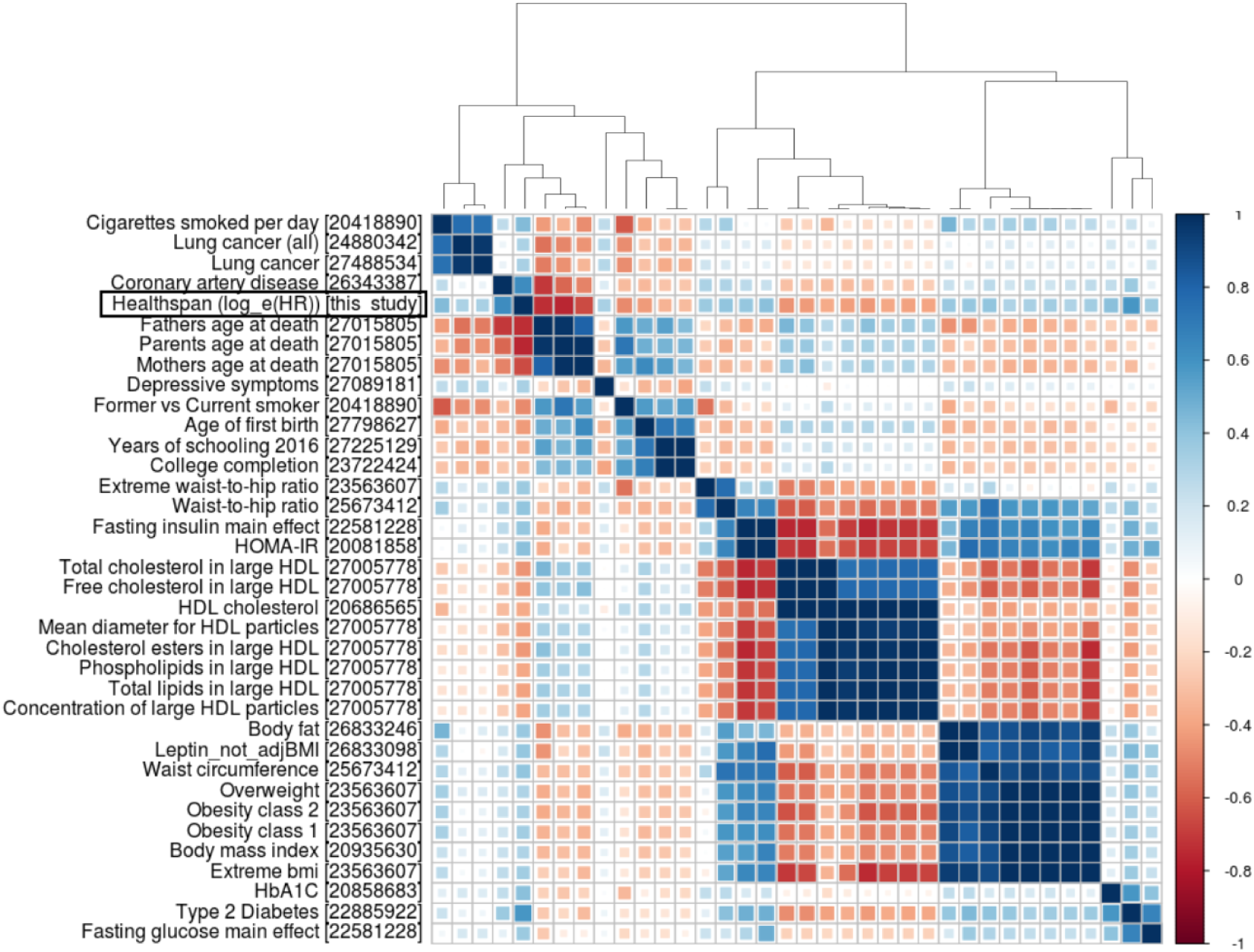
Heatmap for 35 traits with strongest genetic correlations with healthspan (|*r_g_*| ≥ 0.3; *p* ≤ 4.3 × 10^−5^). PMID references are placed in square brackets. Note the absence of genetic correlation between the healthspan and the Alzheimer disease traits (*r_g_* = −0.03).

### E. Functional annotation *in*-*silico*

For the five replicated loci we selected SNPs that most likely include the functional variant (99% credible set). In total, we picked 83 SNPs (Table S5) for further variant effect predictor analysis. The results of variant effect predictor [24] annotation are presented in Table S6. We observed that for a locus on chromosome 6 at 161Mb rs3798220, the variant was missense for two transcripts of the *LPA* gene; for both transcripts rs3798220 was classified as probably damaging by PolyPhen (although it was classified as a tolerated variant by SIFT).

DEPICT [25, 26] analysis using first the 14 SNPs from Supplementary Table 1, and then a larger set of 135 independent SNPs with *p* ≤ 10^−5^ (Table S4) did not yield any significant gene-sets or tissues/cells types enrichment, or prioritized genes (*FDR* > 0.2, Tables S3 and S4).

Finally, we investigated the overlap between associations obtained here and elsewhere, using the phenoscaner v1.1 database [27]. For the twelve most significant SNPs (Table I) we looked up traits that have demonstrated genome-wide significant (*p* < 5 × 10^−8^) associations at the same or at strongly (*r*^2^ < 0.8) linked SNPs. The results are summarized in Table S8. For the five replicated loci we observed co-associations with a number of complex traits. The loci on chromosome 2 at 202 Mb (nearest gene *ALS2CR12*) associated with melanoma skin cancer [28] and esophageal squamous cell carcinoma [29]. Next, loci on chromosome 6 at 0.4 Mb (*IRF4*) associated with different aspects of pigmentation, such as color of skin, eye and hair, pigmentation, tanning and freckles [30, 31], but also with non-melanoma skin cancer [31] and the mole count in cutaneous malignant melanoma families [32]. Two loci (on chromosome 6 at 161 Mb and on chromosome 9 at 22 Mb, *LPA* and *CDKN2B*-*AS1* respectively) were associated with coronary artery disease, myocardial infarction, LDL and cholesterol levels [16, 33]. The remaining replicated locus on chromosome 10 at 114 Mb (*TCF7L2*) was associated with glucose levels, BMI and type 2 diabetes [34, 35].

### F. Effects of known lifespan-associated loci onto healthspan

We have compared whether SNPs, previously reported to be associated with lifespan and (extreme) longevity are associated with healthspan (Table S13) in our data. Interestingly, we observed a very strong enrichment in low p-values (of the 16 tested SNPs, nine had *p* < 0.1), although it should be noted that some SNPs we tested fall into the same region and some were discovered using the same resource (UKB). After correction for multiple testing, we find that four variants (located in or near *CDKN2B*, *ABO*, *LPA*, and *HLA*-*DQA1*), which have been reported to be associated with (extreme) longevity were also significantly associated with the healthspan; two of these variants reached genome-wide significance and were independently discovered as healthspan loci in this study. For the four loci, the allele that was previously reported as being associated with increased longevity, was the one associated with increased healthspan in our data.

## III. DISCUSSION

Risks of death associated with specific age-related diseases increase exponentially at similar rates irrespective of the conditions. The prevalence of disease and therefore the frailty index are also exponential functions of age with the doubling time matching approximately the mortality rate doubling time, see, e.g., [36, 37]. In this study, we observed that the incidence of the diseases of age also growths exponentially with age at nearly the same rates. The manifestly close relation between the diseases and mortality suggests that the healthspan may be a very relevant aging phenotype: the first morbidity signifies the end of the functional or disease-free period, the healthspan, and may signal a transition into a biologically or clinically distinct and relatively short-lived state, linked with the progressive accumulation of frailty, multimorbidity, and death.

Since the first morbidity risk grows exponentially with age, we proposed to employ the probabilistic language of Cox-Gompertz proportional models to test for associations between the demographic and genetic variables, on the one hand, and healthspan, on the other. For example, the Cox-Gompertz model hazard ratio associated with male sex gives 2.5 years for the estimated healthspan difference between the males and females. The Cox-Gompertz lifespan difference of 3.2 years for the same cohort. Although females in UK (the population relevant to this study) live longer than males, the gap between the sexes has decreased over time and is now 3.6 years [38]. The number is very close to our healthspan difference estimate. It is therefore intriguing to see if this numerical coincidence is the model artifact, or if indeed the observed difference in the lifespans could be entirely attributed to the difference in healthspan.

Since gene variant contributions to health- and lifespan are usually small, we obtain the corresponding effect size and statistics estimates with the help of a simple perturbative procedure first proposed in [39] and adopted here. It resembles a regression of the independent variable (the gene variant, in our case) against the Martingale residual of the proportional hazard model, the difference between the predicted and the observed morbidity, see, e.g., [7]. We obtained explicit analytic expressions for the regression coefficient and statistics for the specific case of parametric Cox-Gompertz mortality model, see Eqs. (A2) and (A3). We suggest using the proposed equations or the relevant generalizations for non-parametric risk models for fast and accurate statistical analysis involving small survival effects.

It is tempting to consider the results of our GWAS as informing about potential anti-aging targets. The healthspan, as well as lifespan, however, is an integrated quantity and therefore may depend on the gene activation patterns during subsequent development stages and/or associated with life-long exposure. Therefore, our GWAS ‘hits’ may not necessarily be good targets for an intervention at advanced ages. The appearance of significant genetic correlations with such traits as the years of schooling and the age of the first birth could be indicators of such possibilities. One possible way to deconvolute the effects of human development, diseases and longevity could thus involve using longitudinal clinical data to see if there are gene variants responsible for the rate of aging or biological aging acceleration separately in every age group to negate the effects of accumulation in the course of development.

The strongest genetic correlate of the healthspan is parental longevity. More specifically, *HLA*-*DQB1*, *LPA* and *CDKN2B* loci identified in relation to healthspan in this study were recently associated with parental longevity, a proxy for lifespan, in [8]. Such overall correlation and specific overlap is indeed a desired property of a longevity-associated or a longevity-proxy phenotype. Other traits, belonging to the same cluster, are firstly coronary artery disease, and then lung cancer, smoking behaviour, age of first birth, and years of schooling (Fig.2). The remaining large clusters correspond to traits associated with diabetes type 2, obesity and lipid metabolism, most of which are known to relate to biological age acceleration, see, e.g., [40]. The findings thus provide further evidence suggesting that healthspan and the related diseases could be controlled by common and highly evolutionary conserved mechanisms, such as nutrient sensing and insulin signalling, most robustly implicated in longevity studies in model animals [1, 41].

The notable absence in our study of the gene variants around the *APOE/TOMM40* locus known for association with early onset of Alzheimer disease [42] requires special consideration. First, as shown in Fig. 1, dementia occurs later in life and its incidence rate appears to grow faster than that of the other diseases investigated here in relation with healthspan. The estimated risk doubling time is shorter and is closer to 5 years, in agreement with, e.g., [43]. Next, we performed the dementia GWAS in the same UKB cohorts and failed to produce strong genetic correlations with the healthspan (Fig. 4; note, however, the appreciable correlation between the dementia and mortality traits). We also note the absence of significant genetic correlations between our healthspan and the LD-hub Alzheimer disease traits (Fig. 5). These findings could be an artifact of the age composition of our discovery cohort and possible under-representation of dementia incidence and its influence on healthspan. It could be, however, an indication of distinct underlying biology between the late life neurodegenerative conditions and the more prevalent diseases of aging, mostly occurring at the earlier age, corresponding to the average lifespan in the population.

Using healthspan as a longevity proxy as exemplified here introduces a new way to investigate interactions among gene variants and phenotypic variation due to effects of lifestyles and living conditions. On a population level, such life-history variables produce a very significant contribution to longevity [44], are inherently stochastic and may not be readily estimated for inclusion in most forms of genetic studies. It would be rather difficult, for example, to extrapolate social status, sleep patterns or food habits across generations for GWAS against parental age at death. Modern large population studies involve prospective cohorts and produce a very rich characterization of the participants, yet at the expense of limited follow-up times and an insufficient number of recorded death or morbidity events. The healthspan as the target phenotype should help bring the best of the two worlds and thus improve the statistical power of longevity GWAS and eventually assist in the discovery of many more genes implicated in the control of human aging and diseases.

The burden of diseases increases with age, and the first morbidity is usually quickly followed by the second and more. Therefore it is worthwhile to understand if the same or different genes than those regulating the onset of the first morbidity (the end of healthspan, as defined in this study) also control the dynamics of multiple morbidities later down the road. The comparison and better understanding of the results of such studies will help to differentiate the biology of health- and life-span. Human development and aging is a multi-stage process, and therefore longevity emerges as a genuinely complex trait. The presented study highlights a need for further systematic advances in aging GWAS methodology to elucidate the practical potential of genetics in diagnosis of aging and, subsequently, help to shape the anti-aging therapeutic target space.

## IV. ACKNOWLEDGEMENT

The work of SSh was supported by Russian Ministry of Science and Education under 5-100 Excellence Programme. The work of YT was supported by the Federal Agency of Scientific Organizations via the Institute of Cytology and Genetics (project #0324-2018-0017). This research has been conducted using the UK Biobank Resource. The study has been funded by Gero LLC.

## Appendix A: Methods

### 1. UK Biobank

UK Biobank is a prospective cohort study of over 500,000 individuals from across the United Kingdom [45]. Participants, aged between 40 and 69, were invited to one of 22 centres across the UK between 2006 and 2010. Blood, urine and saliva samples were collected, physical measurements were taken, and each individual answered an extensive questionnaire focused on questions of health and lifestyle. All participants gave written informed consent and the study was approved by the North West Multicentre Research Ethics Committee. UKB has Human Tissue Authority research tissue bank approval, meaning separate ethical approvals are not required to use the existing data. UKB provided genotyping information for 488,377 individuals. Data access to UKB was granted under MAF 21988. Phenotypes and genotypes were downloaded directly from UKB.

### 2. Genotyping and imputations

UKB participants were genotyped on two slightly different arrays and quality control was performed by UKB [46]. 49,950 samples were genotyped as part of the UK BiLEVE study using a newly designed array, with 438,427 remaining samples genotyped on an updated version (UK Biobank Axiom array), both manufactured by Affymetrix (96% of SNPs overlap between the arrays). Samples were processed and genotyped in batches approx. 5000 samples each. In brief, SNPs or samples with high missingness, multi-allelic SNPs and SNPs with batchwise departures from HardyWeinberg equilibrium were removed from the data set. After quality control, genotypes were available for 488k subjects at 805k sites. UKB provided 40 principal components of genetic relatedness (UKB field id 22009) and a binary assessment of whether subjects were genetically Caucasian (UKB field id 22006), based on principal components analysis of their genetic data.

Imputed data were prepared by UKB. In summary, autosomal phasing was carried out using a version of SHAPEIT2 [47] modified to allow for very large sample sizes. Imputation was carried out using IMPUTE2 [48] using the merged UK10K and 1,000 Genomes Phase 3 reference panels to yield higher imputation accuracy of British haplotypes. The imputations resulted in 92,693,895 SNPs, short indels and large structural variants, imputed in 488,377 individuals (UK-Biobank. Genotype Imputation and Genetic Association Studies of UK Biobank.

### 3. Discovery and replication samples

For the discovery and replication we used only the data from PCA cohort (QC passed, Data-Field 22020, N = 407,208). For the discovery set we selected 300,447 British individuals of European ancestry (EA) according to the genetic principal components provided by the UK Biobank who were not included in UK BiLEVE study (UKB Resource 531). For replication, we used a combination of the UK Biobank participants not included in the discovery set that comprised rest of EA individuals (self-reported white, data-field 21000, n = 81,099), individuals of African ancestry (self-reported Africans, n = 3,073), individuals of South Asian ancestry (Indian, Pakistani, and Bangladeshi; n = 6,921), Chinese individuals (n = 1,422) and Caribbean individuals (n=3,799). Remaining self-declared ethnicities that were mixed, or were ambiguous (Other ethnic group, Prefer not to answer, Not available) were not analysed. To reduce the risk of bias due to population stratification, all groups were analyzed separately followed by a meta-analysis. Total resulting sample size for replication was 96,313 individuals. For more details see Supplementary Table 8.

The replication threshold was set as *p* < 0.05/12 = 0.004. For each SNP, statistical power (or probability) of replication was estimated using the fact that under alternative hypothesis (*H*1 : *β* ≠ 0) the test statistics *T*^2^ from replication sample is expected to follow the
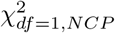
distribution, where *NCP* is the expected non-centrality parameter computed as
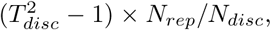
where
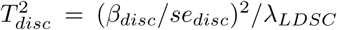
is test statistic for particular SNP in discovery cohort, corrected for LD score regression interecept λ_*LDSC*_, *N_rep_* is the sample size of the replication cohort and *N_disc_* is the sample size of the discovery cohort. The the power of replication is equal to the probability that such distributed statistics would exceed the threshold value *k* = 8.2 that corresponds to right-hand integral of 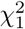
equal to 0.004.

### 4. Incidence of diseases calculation from UKB data

We used in-patient hospital admissions data (UKB data category 2000) and self-reported diagnoses obtained via verbal interview (UKB data category 100074) to extract information in relation to the disease history, the nature of and the age at the available diagnosis. For each of the condition, we follow the instructions similar to the ones given by the UK Biobank outcome adjudication group for algorithmic-defined stroke and MI (UKB data category 42). For each selected condition, except for cancer and death we compile a list of hospital data codes (ICD-10, Supplementary Table 11) and self-reported data codes (UKB data coding 6) that defines these conditions in our study. We used National cancer registries linkage to UKB (UKB data category 100092) in addition to hospital data for cancer and National death registries linkage to UKB (UKB data category 100093) to define death event. First, for each condition we set the age of first occurrence of any of corresponding hospital data codes as age this condition was manifested. Next, if there was missing hospital data (for hospital data it is impossible to distinguish between missing data and absence of any disease) we added self-reported data if there was any. Therefore we obtained age each condition was occurred. The minimal age from this data set for every individual from UKB was taken as age the healthspan terminates.

By definition, the incidence rate of a disease is the limit *m*(*t*) = Δ*t*^−1^*N_d_*(*t*, Δ*t*)/*N_h_*(*t*) when Δ*t* is sufficiently small. Here *t* is the age, *N_h_*(*t*) is the number of people healthy at the age *t* and *N_d_*(*t*, Δ*t*) is the number of people diagnosed between the ages *t* and *t* + Δ*t* (both *N_h_* and *N_d_* are presumed to be large). This definition does not rely on any specific underlying model. In practice, datasets are of limited size and the interval Δ*t* cannot be made arbitrarily small, and therefore precautions should be taken to avoid possible artifacts in the calculation. To compute the incidence rate at a given age *t*, one shall consider a set of participants ϒ(*t*, Δ*t*) defined as those who are healthy at the age *t* and whose health status is available in the whole age range [*t*,*t* + Δ*t*): ϒ(*t*, Δ*t*) = {*u*|((*δ^u^* =0) ∨ (*δ_u_* = 1 ∧ *t* ≥
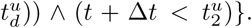
Here *u* is the participant’s id, *δ^U^* = 1 if the participant was diagnosed and *δ^u^* = 0 otherwise,
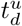
is the age when diagnosed, and
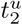
is the maximal age at which the information about the diagnosis (if any) would still be recorded. From this *N_h_*(*t*) = |ϒ(*t*, Δ*t*)| and *N_d_*(*t*, Δ*t*) = |{*u* ∈ ϒ(*t*, Δ*t*)|*δ^U^* =
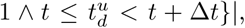
where |..| is the size of the set.

The maximum follow-up age
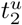
does not coincide with the age at the diagnosis
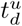
and shall be inferred from the study setup. Assuming
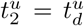
for diagnosed participants would overestimate the risks. Also, the age is often rounded and hence Δ*t* may be not large enough to treat the rounding errors as negligible. We addressed the issue by consistently using half-open intervals [..) definitions. Finally, our prescription relies on the implicit assumption, that the diagnosis does not influence the enrolment. This is not always true. If someone is dead, this would, naturally, prevent that person from being enrolled at a greater age. This can be addressed by the following modification:
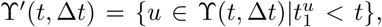
where
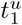
is the age at enrollment. In this study, we assumed that the enrolment in UKB was not biased by diagnoses and thus we used the ϒ for all diseases and conditions, ϒ′ participants set was only employed for the mortality rate calculation.

### 5. Cox-Gompertz proportional hazards model and healthspan

By design of the UKB study, every participant is admitted into the cohort at the age
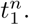
According to the medical history information, the participant may be diagnosed with any of the diseases relevant to determination of lifespan at the age of the first
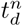
(if applicable). By the end of the followup age,
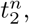
we labeled every study participant as frail, *δ^n^* = 1, if the participant is already diagnosed with any of the diseases,
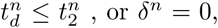
otherwise.

Under then Cox-Gompertz proportional hazards model the risks of frailty acquisition or healthspan end at the age *t* is *h*(*t*, *x^n^*) = *h*_0_ exp(Γ*t* + *βx^n^*), where *x^n^* is a vector of age-independent parameters, characterizing the participant. Here *h*_0_, Γ, and *β* are the baseline morbidity incidence, the Gompertz exponent and the log-odds-ratio regression coefficients vector, the model parameters. The (negative log of) likelihood of the data can be presented in the following form:

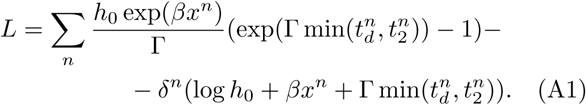

Given a necessary amount of data the model parameters could be obtained by the likelihood maximization or, equivalently, minimization of the cost function *L*.

We built the first version of the Cox-Gompertz healthspan model by including UKB participants information, including gender and the first genetic principal components variables, assessment center codes and genotyping batch labels (see supplementary Table 14 for the summary of the model parameters). The morbidity incidence growth rate is 0.098 per year, which corresponds to a doubling time of seven years, compatible with the mortality rate doubling time of approximately 8 from the Gompertz mortality law. As expected, being male is a risk factor (log-hazard ratio, log(*HR*) = 0.26 at the significance level of *p* ≪ 10^−10^) corresponding to an average healthspan difference of about five years. The genetic principal components PC4 and PC5 are also highly significant (log(*HR*) ≈ 3 × 10^−2^, *p* ≪ 10^−10^). The average healthspan or lifespan can be estimated from Cox-Gompertz model parameters as *t* ≈ (ln(Γ/*h*_0_) − *γ*)/Γ, where *γ* = 0.577 is the Euler-Mascheroni constant, see, e.g., [49].

### 6. Gene variant-healthspan association testing

If the participants state vector
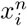
is extended by the genetic variants variables *s^n^*, in principle, the model has to be re-evaluated, to obtain a new versions of all model parameters. We do not expect, however, large effects of any of the gene variants on lifespan. Therefore the model parameters should not change much as well and the variation of the Cox-Gompertz model with respect to the genetic variables can be accurately obtained by iterations, using the model from A 5 as the zeroth order approximation (see a related example of a perturbation theory application in a proportional hazards model involving prediction of all-cause mortality in [39]).

We note, however, that the simultaneous determination of the weak effects of a gene on the baseline hazard *h*_0_ and the rate of aging Γ is an ill-defined mathematical problem [49]. Only the combination of the two parameters, the change in the life expectancy can be determined with accuracy. We therefore fix the Gompertz exponent Γ to its most probable value in the zeroth order model and allow for all other model parameters adjustment. The perturbation theory expansion for the small effect *β_s_* associated with the gene variants yields (the derivation is not shown):

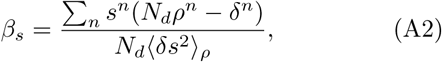

where, for convenience, we introduced the weights

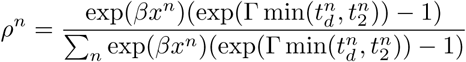

normalized in such a way that ∑_*n*_ *ρ_n_* = 1. We used the notation 〈*δs*^2^〉_*ρ*_ for the corresponding weighted average.

The effect determination error

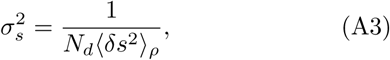

and hence the statistical power of the gene variant association with the healthspan is explicitly dependent on the number of people with diagnoses, *N_d_* = ∑_*n*_ *δ^n^*.

In our analyses, we used imputed variants with the expected effective minor allele count (defined as twice the minor allele frequency multiplied by sample size and by the imputation quality) more than 200 for discovery cohort genotypes and imputation info score (as IMPUTE info, calculated by RegScan for discovery cohort with `--info2` option) more than 0.7.

### 7. Conditional and joint multi-SNP analysis

Conditional and joint analysis (COJO) as implemented in the program GCTA [50] was used to find SNPs independently associated with the phenotypes of interest. As input, this method uses (meta-analysis) summary statistics and a reference sample that is utilised for the LD estimation. The method starts with the “top SNP” (the one with smallest p-value, conditional that *p* < *p*_0_, where *p*_0_ is specific threshold defined by user) as provided by the summary-level data and then the p-values for all the remaining SNPs are calculated conditional on the selected SNP. The algorithm then selects the next top SNP in the conditional analysis (provided *p* < *p*_0_) and proceeds to fit all the selected SNPs in the model dropping all those SNPs with p-values > *p*_0_. The iteration continues until no SNP is added or dropped from the model thus finding a subset of associated SNPs with a threshold for LD (*r*^2^ < 0.9) among SNPs. Finally, a joint analysis of the subset of associated SNPs is performed. We had performed analyses with *p*_0_ = 5 × 10^−8^ and *p*_0_ = 1 × 10^−5^.

As the LD reference, we used a sub-sample of 10,000 people, randomly chosen from the total set of 120,286 people used for GWAS discovery phase. Additional to our previous SNP filters described in the “Association testing” section, in selecting LD reference data, we further filtered out the SNPs with imputation info scores less than 0.7 and minor allele frequencies (MAF) less than 0.002.

### 8. Heritability and genetic correlation analyses

We used LD hub and ldsc [51] tools for estimation of captured heritability and genetic correlations between HS and different traits and common diseases [51]. A total of 231 traits were analyzed after removing duplicates via using only the most recent study for each trait as indicated by the largest PMID number. Genetic correlations between HS and the traits with p-value 4.3e-5 (Bonferroni corrected, 0.01/231) were considered statistically significant. Pair-wise genetic correlations between all the traits selected as described above were obtained from the LD-hub. To focus on the largest magnitude genetic correlations, we selected only the traits with absolute values of genetic correlations with HS more than 0.3. This filtering led to the total of 36 traits (including HS). Clustering and visualization was carried out using corrplot package for R and basic hclust function. For clustering, we estimated squared Euclidean distances by subtracting absolute values of genetic correlation from 1 and used Ward’s clustering method.

For genetic correlation anasysis between each disease comprising healthspan phenotype and healthspan itself we used LDSC (LD Score) v1.0.0 software. Genotype calls were filtered by *MAF* > 0.01 using LDSC ‘munge-sumstats’ script to produce total 659,079 variants valid for downstream analysis. Genomic reference was constructing by randomly sampling 10,000 individuals from the UKB population. Then, we ran LDSC genetics correlation analysis with default parameters and input data as described above. Cross-correlations can be seen at figure 4 and Table S15.

For analysis of heritability, genomic control inflation factor λ [13] and genetics correlations we have used SNPs defined by overlap between our set of SNPs and ‘high quality SNPs’ as suggested by the authors of the LD hub (these represent common HapMap3 SNPs that usually have high imputation quality; also, this set excludes HLA region) [14], 1,162,742 SNPs in total).

### 9. In silico functional analysis

#### Variant effect prediction (VEP)

We used PAINTOR software to prepare the set of SNPs for functional annotation. For this analysis, we provided PAINTOR with clumping results, LD matrices and annotation files. Using PLINK [52] and 10,000 samples reference set described above (the same subset as used in COJO and DEPICT analyses) we performed clumping analysis with ‘p1’ and ‘p2’ p-value threshold parameters set to 5 × 10^−8^, ‘r2’ set to 0.1 and MAF¿0.002. Then, using the same reference set we generated pair-wise correlation matrix for all SNPs in each region in clumping analysis results using plink `--r` option. When running PAINTOR, we did not use annotations; we changed options controlling input and output files format only, and otherwise we have used default parameters. In the next step, all output results were aggregated into one file and SNPs marked by PAINTOR as 99% credible set we chosen for functional annotation by VEP with GRCH37 genomic reference.

#### Gene-set and tissue/cell enrichment analysis

For prioritising genes in associated regions, gene set enrichment and tissue/cell type enrichment analyses, we have used the DEPICT software v. 1 rel. 194 [25] with following parameters: flagdoci = 1; flag_genes = 1; flag_genesets = 1; flag_tissues = 1; param_ncores = 10. Independent (as selected by COJO procedure) variants with *p* < 5 × 10^−8^ (14 SNPs) and *p* < 10^−5^ (135 SNPs) has resulted from these analyses. We have used UKB subset of 10,000 individuals for computations of LD (the same subset as used for COJO analysis).

#### Pleiotropy with complex traits

We investigated the overlap between associations obtained here and elsewhere, using PhenoScaner v1.1 database. For five replicated SNPs (Table 1) we looked up traits that have demonstrated genome-wide significant (*p* < 5 × 10^8^) association at the same or at strongly (*r*2 < 0.8) linked SNPs.

## Appendix B: Supplementary info

### 1. Supplementary information legend

- Supplementary Table 1: Supplementary Table 1. Extended results for variants, tagging regions, significantly associated with lifespan in the discovery sample of 300,447 individuals; association between these variants and lifespan in the replication sample of 55,276 self-reported British individuals and 41,037 individuals of other ethnicities (total *N* = 96, 313).
- Supplementary Table 2: Genetic correlations between frailty and different traits, as estimated by LD score regression.
- Supplementary Table 3A: Gene set enrichment analysis with DEPICT (input SNPs with p¡5e-8).
- Supplementary Table 3B: Gene prioritisation with DEPICT (input SNPs with p¡5e-8).
- Supplementary Table 3C: Tissue enrichment analysis with DEPICT (input SNPs with p¡5e-8).
- Supplementary Table 4A: Gene set enrichment analysis with DEPICT (input SNPs with p¡1e-5).
- Supplementary Table 4B: Gene prioritisation with DEPICT (input SNPs with p¡1e-5).
- Supplementary Table 4C: Tissue enrichment analysis with DEPICT (input SNPs with p¡1e-5).
- Supplementary Table 5: 99% credible set for SNPs implicated as genome-wide significant and independent by COJO analysis.
- Supplementary Table 6: Summary from analysis of variants from 99% credible set with variant effect predictor (VEP).
- Supplementary Table 7: Replication cohort composition.
- Supplementary Table 8A: Other traits associated to the regions showing significant association with predicted frailty (GWAS).
- Supplementary Table 8B: Other traits associated to the regions showing significant association with predicted frailty (eQTL).
- Supplementary Table 8C: Other traits associated to the regions showing significant association with predicted frailty (Metabolites).
- Supplementary Table 9: All significant SNPs.
- Supplementary Table 10: Cox-Gomperz summary statistics for single diseases
- Supplementary Table 11. Disease codes for healthspan composition.
- Supplementary Table 12.
- Supplementary Table 13. Variants previously implicated in studies of longevity and aging.
- Supplementary Table 14. Cox-Gompertz model parameters estimation.

## 2. Authors contributions

Analysis implementation: AZ, YT, SS, G; Experimental design: YA, PF, LM; Manuscript writing: AZ, YT, SS, G, LM, YA, PF.

## References

[1] T. Niccoli and L. Partridge, Current Biology 22, R741 (2012).

[2] S. L. Andersen, P. Sebastiani, D. A. Dworkis, L. Feldman, and T. T. Perls, Journals of Gerontology Series A: Biomedical Sciences and Medical Sciences 67, 395 (2012).

[3] B. K. Kennedy, S. L. Berger, A. Brunet, J. Campisi, A. M. Cuervo, E. S. Epel, C. Franceschi, G. J. Lithgow, R. I. Morimoto, J. E. Pessin, et al., Cell 159, 709 (2014).

[4] J. Deelen, M. Beekman, H.-W. Uh, L. Broer, K. L. Ayers, Q. Tan, Y. Kamatani, A. M. Bennet, R. Tamm, S. Trompet, et al., Human molecular genetics 23, 4420 (2014).

[5] K. Fortney, E. Dobriban, P. Garagnani, C. Pirazzini, D. Monti, D. Mari, G. Atzmon, N. Barzilai, C. Franceschi, A. B. Owen, et al., PLoS Genet 11, e1005728 (2015).

[6] Y. Zeng, C. Nie, J. Min, X. Liu, M. Li, H. Chen, H. Xu, M. Wang, T. Ni, Y. Li, et al., Scientific reports 6, 21243 (2016).

[7] P. K. Joshi, K. Fischer, K. E. Schraut, H. Campbell, T. Esko, and J. F. Wilson, Nature communications 7, 11174 (2016).

[8] P. K. Joshi, N. Pirastu, K. A. Kentistou, K. Fischer, E. Hofer, K. E. Schraut, D. W. Clark, T. Nutile, C. L. Barnes, P. R. Timmers, et al., Nature communications 8, 910 (2017).

[9] A. F. McDaid, P. K. Joshi, E. Porcu, A. Komljenovic, H. Li, V. Sorrentino, M. Litovchenko, R. P. Bevers, S. Rüeger, A. Reymond, et al., Nature communications 8, 15842 (2017).

[10] B. Gompertz, Philosophical transactions of the Royal Society of London 115, 513 (1825).

[11] W. M. Makeham, The Assurance Magazine and Journal of the Institute of Actuaries 8, 301 (1860).

[12] N. A. Tutkun and H. Demirhan, Hacettepe Journal of Mathematics and Statistics 45, 1621 (2016).

[13] B. Devlin and K. Roeder, Biometrics 55, 997 (1999).

[14] B. K. Bulik-Sullivan, P.-R. Loh, H. K. Finucane, S. Ripke, J. Yang, N. Patterson, M. J. Daly, A. L. Price, B. M. Neale, S. W. G. of the Psychiatric Genomics Consortium, et al., Nature genetics 47, 291 (2015).

[15] J. Zheng, A. M. Erzurumluoglu, B. L. Elsworth, J. P. Kemp, L. Howe, P. C. Haycock, G. Hemani, K. Tansey, C. Laurin, B. S. Pourcain, et al., Bioinformatics 33, 272 (2017).

[16] M. Nikpay, A. Goel, H.-H. Won, L. M. Hall, C. Willenborg, S. Kanoni, D. Saleheen, T. Kyriakou, C. P. Nelson, J. C. Hopewell, et al., Nature genetics 47, 1121 (2015).

[17] A. P. Morris, B. F. Voight, T. M. Teslovich, T. Ferreira, A. V. Segre, V. Steinthorsdottir, R. J. Strawbridge, H. Khan, H. Grallert, A. Mahajan, et al., Nature genetics 44, 981 (2012).

[18] N. Soranzo, S. Sanna, E. Wheeler, C. Gieger, D. Radke, J. Dupuis, N. Bouatia-Naji, C. Langenberg, I. Prokopenko, E. Stolerman, et al., Diabetes 59, 3229 (2010).

[19] H. Furberg, Y. Kim, J. Dackor, E. Boerwinkle, N. Franceschini, D. Ardissino, L. Bernardinelli, P. M. Mannucci, F. Mauri, P. A. Merlini, et al., Nature genetics 42, 441 (2010).

[20] J. Dupuis, C. Langenberg, I. Prokopenko, R. Saxena, N. Soranzo, A. U. Jackson, E. Wheeler, N. L. Glazer, N. Bouatia-Naji, A. L. Gloyn, et al., Nature genetics 42, 105 (2010).

[21] N. Barban, R. Jansen, R. De Vlaming, A. Vaez, J. J. Mandemakers, F. C. Tropf, X. Shen, J. F. Wilson, D. I. Chasman, I. M. Nolte, et al., Nature genetics 48, 1462 (2016).

[22] L. C. Pilling, J. L. Atkins, K. Bowman, S. E. Jones, J. Tyrrell, R. N. Beaumont, K. S. Ruth, M. A. Tuke, H. Yaghootkar, A. R. Wood, et al., Aging (Albany NY) 8, 547 (2016).

[23] J. Kettunen, A. Demirkan, P. Würtz, H. H. Draisma, T. Haller, R. Rawal, A. Vaarhorst, A. J. Kangas, L.-P. Lyytikainen, M. Pirinen, et al., Nature communications 7, 11122 (2016).

[24] W. McLaren, L. Gil, S. E. Hunt, H. S. Riat, G. R. Ritchie, A. Thormann, P. Flicek, and F. Cunningham, Genome biology 17, 122 (2016).

[25] T. H. Pers, J. M. Karjalainen, Y. Chan, H.-J. Westra, A. R. Wood, J. Yang, J. C. Lui, S. Vedantam, S. Gustafsson, T. Esko, et al., Nature communications 6, 5890 (2015).

[26] G. P. Consortium et al., Nature 467, 1061 (2010).

[27] J. R. Staley, J. Blackshaw, M. A. Kamat, S. Ellis, P. Surendran, B. B. Sun, D. S. Paul, D. Freitag, S. Burgess, J. Danesh, et al., Bioinformatics 32, 3207 (2016).

[28] J. H. Barrett, M. M. Iles, M. Harland, J. C. Taylor, J. F. Aitken, P. A. Andresen, L. A. Akslen, B. K. Armstrong, M.-F. Avril, E. Azizi, et al., Nature genetics 43, 1108 (2011).

[29] C. C. Abnet, Z. Wang, X. Song, N. Hu, F.-Y. Zhou, N. D. Freedman, X.-M. Li, K. Yu, X.-O. Shu, J.-M. Yuan, et al., Human molecular genetics 21, 2132 (2012).

[30] N. Eriksson, J. M. Macpherson, J. Y. Tung, L. S. Hon, B. Naughton, S. Saxonov, L. Avey, A. Wojcicki, I. Pe’er, and J. Mountain, PLoS genetics 6, e1000993 (2010).

[31] M. Zhang, F. Song, L. Liang, H. Nan, J. Zhang, H. Liu, L.-E. Wang, Q. Wei, J. E. Lee, C. I. Amos, et al., Human molecular genetics 22, 2948 (2013).

[32] D. L. Duffy, M. M. Iles, D. Glass, G. Zhu, J. H. Barrett, V. Höiom, Z. Z. Zhao, R. A. Sturm, N. Soranzo, C. Hammond, et al., The American Journal of Human Genetics 87, 6 (2010).

[33] C. J. Willer, E. M. Schmidt, S. Sengupta, G. M. Peloso, S. Gustafsson, S. Kanoni, A. Ganna, J. Chen, M. L. Buchkovich, S. Mora, et al., Nature genetics 45, 1274 (2013).

[34] K. J. Gaulton, T. Ferreira, Y. Lee, A. Raimondo, R. Mägi, M. E. Reschen, A. Mahajan, A. Locke, N. W. Rayner, N. Robertson, et al., Nature genetics 47, 1415 (2015).

[35] D. Shungin, T. W. Winkler, D. C. Croteau-Chonka, T. Ferreira, A. E. Locke, R. Mägi, R. J. Strawbridge, T. H. Pers, K. Fischer, A. E. Justice, et al., Nature 518, 187 (2015).

[36] R. Yu, W.-C. Wu, J. Leung, S. C. Hu, and J. Woo, International journal of environmental research and public health 14, 1096 (2017).

[37] A. Mitnitski and K. Rockwood, Biogerontology 17, 199 (2016).

[38] “National life tables, uk: 2014 to 2016,” (2017).

[39] T. V. Pyrkov, K. Slipensky, M. Barg, A. Kondrashin, B. Zhurov, A. Zenin, M. Pyatnitskiy, L. Menshikov, S. Markov, and P. O. Fedichev, Scientific Reports 8, 5210 (2018).

[40] S. Horvath, W. Erhart, M. Brosch, O. Ammerpohl, W. von Schönfels, M. Ahrens, N. Heits, J. T. Bell, P.-C. Tsai, T. D. Spector, et al., Proceedings of the National Academy of Sciences 111, 15538 (2014).

[41] M. Piper, C. Selman, J. McElwee, and L. Partridge, Journal of internal medicine 263, 179 (2008).

[42] A. Roses, M. Lutz, H. Amrine-Madsen, A. Saunders, D. Crenshaw, S. Sundseth, M. Huentelman, K. Welsh-Bohmer, and E. Reiman, The pharmacogenomics journal 10, 375 (2010).

[43] K. Ziegler-Graham, R. Brookmeyer, E. Johnson, and H. M. Arrighi, Alzheimer’s & dementia: the journal of the Alzheimer’s Association 4, 316 (2008).

[44] L. Wilhelmsen, K. Svärdsudd, H. Eriksson, A. Rosengren, P.-O. Hansson, C. Welin, A. Odén, and L. Welin, Journal of internal medicine 269, 441 (2011).

[45] C. Sudlow, J. Gallacher, N. Allen, V. Beral, P. Burton, J. Danesh, P. Downey, P. Elliott, J. Green, M. Landray, et al., PLoS medicine 12, e1001779 (2015).

[46] C. Bycroft, C. Freeman, D. Petkova, G. Band, L. T. Elliott, K. Sharp, A. Motyer, D. Vukcevic, O. Delaneau, J. O’Connell, et al., bioRxiv, 166298 (2017).

[47] O. Delaneau, J.-F. Zagury, and J. Marchini, Nature methods 10, 5 (2013).

[48] B. Howie, J. Marchini, and M. Stephens, G3: Genes, Genomes, Genetics 1, 457 (2011).

[49] A. E. Tarkhov, L. I. Menshikov, and P. O. Fedichev, Journal of theoretical biology 416, 180 (2017).

[50] J. Yang, T. Ferreira, A. P. Morris, S. E. Medland, P. A. Madden, A. C. Heath, N. G. Martin, G. W. Montgomery, M. N. Weedon, R. J. Loos, et al., Nature genetics 44, 369 (2012).

[51] B. Bulik-Sullivan, H. K. Finucane, V. Anttila, A. Gusev, F. R. Day, P.-R. Loh, L. Duncan, J. R. Perry, N. Patterson, E. B. Robinson, et al., Nature genetics 47, 1236 (2015).

[52] C. C. Chang, C. C. Chow, L. C. Tellier, S. Vattikuti, S. M. Purcell, and J. J. Lee, Gigascience 4, 7 (2015).

